# ChromPolymerDB: A High-Resolution Database of Single-Cell 3D Chromatin Structures for Functional Genomics

**DOI:** 10.1101/2025.09.27.678816

**Authors:** Min Chen, Lin Du, Siyuan Zhao, Bowei Ye, Pourya Delafrouz, Hammad Farooq, Debaleena Chattopadhyay, G. Elisabeta Marai, Zhifeng Shao, Jie Liang, Daniel M. Czajkowsky, Constantinos Chronis

**Author notes:** To whom correspondence should be addressed.; Correspondence may also be addressed to Daniel M. Czajkowsky. The first two authors should be regarded as Joint First Authors.

## Abstract

The three-dimensional (3D) organization of chromatin plays a critical role in regulating gene expression and genomic processes like DNA replication, repair, and genome stability. Although these processes occur at the individual-cell level, most chromatin structure data are derived from population-averaged assays, such as Hi-C, obscuring the heterogeneity of single-cell conformations. To address this limitation, we developed a polymer physics-based modelling framework, the Sequential Bayesian Inference Framework (sBIF), that deconvolutes bulk Hi-C data to reconstruct single-cell 3D chromatin conformations. To support a broader use of sBIF, we created ChromPolymerDB, a publicly accessible, high-resolution database of single-cell chromatin structures inferred by sBIF. The database contains ∼10^8^ reconstructed 5 kb-resolution single cell structures, spanning over 60,000 genomic loci across 50 human cell types and experimental conditions. ChromPolymerDB features an interactive web interface with tools for 3D structural analysis and multi-omics integration. Users can explore associations between chromatin conformation and gene expression, epigenetic modifications, and regulatory elements. The platform also supports comparative analyses to identify structural changes across cell types, developmental stages, or disease contexts. ChromPolymerDB offers a unique resource for researchers studying the relationship between genome architecture and gene regulation, and for advancing comparative 3D genomics. ChromPolymerDB is available online at https://chrompolymerdb.bme.uic.edu/.

**GRAPHICAL ABSTRACT:** 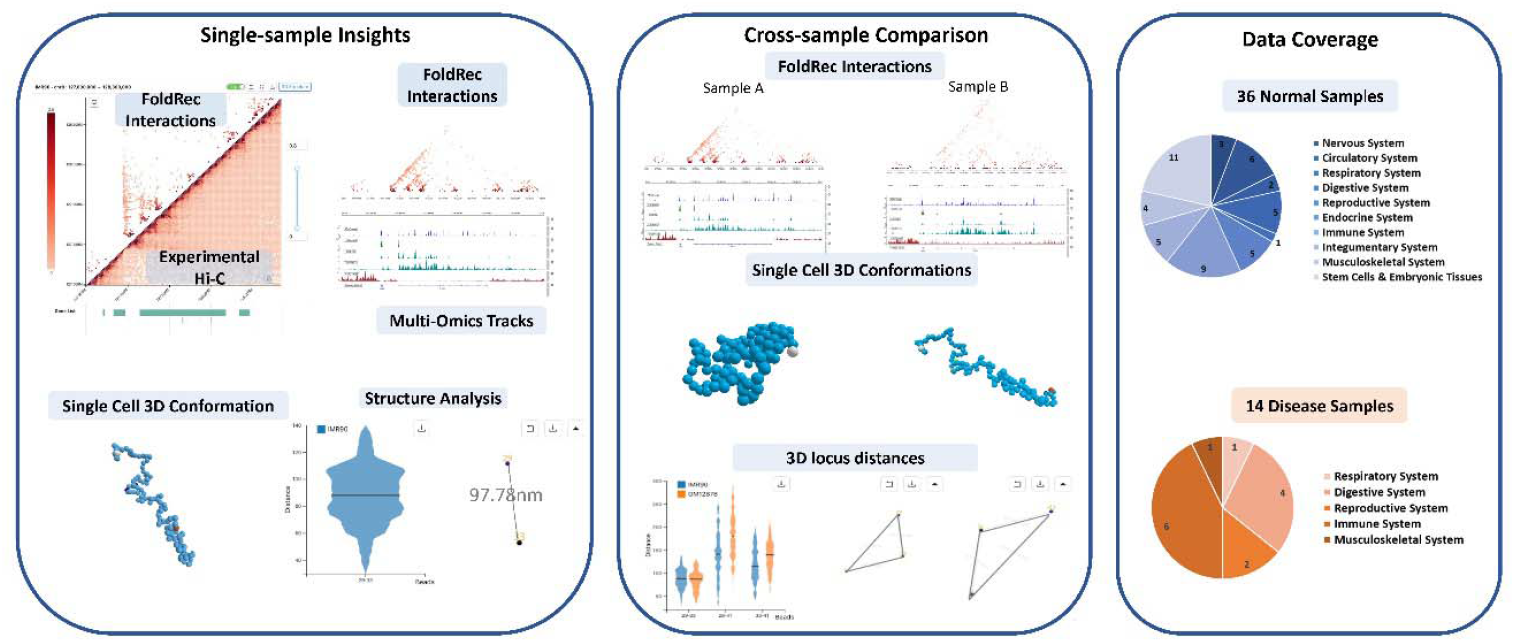

## INTRODUCTION

The three-dimensional (3D) organization of chromatin plays a central role in regulating virtually all genomic processes, including gene expression (1–3), DNA replication, and repair (4). Chromatin architecture underpins the establishment and maintenance of cellular identity and is dynamically remodeled during biological transitions such as differentiation (5–7), development (7–9), and disease progression (10–14). Advances in chromosome conformation capture methods - particularly Hi-C - have enabled genome-wide mapping of chromatin interactions, revealing that genome function is hierarchically organized across length scales, from inter-chromosomal compartments to topologically associating domains (TADs) and fine-scale chromatin loops (15–25).

Despite these insights, most Hi-C datasets are derived from bulk populations of ∼10^6^ cells (16), yielding ensemble-averaged contact maps that obscure cell-to-cell variability (26). Although gene regulation and genome function occur at the level of an individual cell, bulk Hi-C does not resolve the precise chromatin architecture within any one cell (27–30). Imaging studies have confirmed substantial heterogeneity in chromatin conformation across single cells (21, 31), and while single-cell Hi-C methods have emerged to address this limitation (27, 28, 32–34), they remain constrained by extreme data sparsity. The limited number of contacts detected per cell prevents reliable reconstruction of genome-wide 3D structures, particularly at the resolution of TADs and loops (35–38).

To overcome these challenges, we developed the Sequential Bayesian Inference Framework (sBIF), a polymer-physics-based approach for inferring single-cell chromatin conformations from population Hi-C data (39). sBIF employs a deep-sampling strategy with minimal physical assumptions and no adjustable parameters. It has been validated across species, from *Drosophila* to human (23, 39–41) demonstrating its ability to reproduce bulk Hi-C patterns while capturing chromatin heterogeneity at the single-cell level.

To facilitate the broader application of this approach, we established ChromPolymerDB, a high-resolution, open-access database of single-cell 3D chromatin structures reconstructed using sBIF. The resource includes ∼10^8^ structures at 5 kb resolution, spanning over 60,000 genomic regions across 50 human cell types and experimental conditions. ChromPolymerDB offers a user-friendly web interface equipped with built-in tools for interactive 3D visualization, multi-omics integration, and structural analyses (such as 3D locus distances), within individual samples and across samples, as well as com companion off-line analysis code (such as clustering, TAD radius of gyration, and multi-body contacts). By enabling cross-modality comparisons with transcriptomics, epigenomics, and imaging data, the database facilitates the discovery of structural rewiring events, such as enhancer hub formation or loop remodeling, that can occur during cellular transitions or between disparate cell fates. By facilitating access to and analysis of 3D chromatin architectures, we thus anticipate that ChromPolymerDB will prove to be a useful resource for investigations of chromatin-gene regulation relationships and comparative 3D genomics.

## MATERIAL AND METHODS

### Data Collection

To generate ChromPolymerDB, we collected 50 high-quality human Hi-C samples from three major public databases: the 4D Nucleome (4DN) Data Portal (42) (n=19), the ENCODE Portal (43, 44) (n=18), and the Gene Expression Omnibus (GEO) (45) (n=13). To provide data that might be useful to most researchers, the following criteria were used for sample selection: (1) human origin; (2) sufficient sequencing depth to achieve 5 kb resolution; and (3) homogeneous cell populations. In some cases, we merged multiple datasets generated under largely similar experimental conditions to achieve higher resolution. Overall, our dataset includes 36 normal cell types (both primary and cultured cells) and 14 disease-associated cell types, covering 10 of the 11 major human physiological systems (except the urinary system).

### Preprocessing Hi-C Data for Modeling

Following a workflow similar to the original sBIF protocol (39), we applied a four-step pre-processing pipeline to prepare input regions for structural modeling: acquisition of .hic files, TAD boundary identification, TAD-based segmentation of modeling regions, and filtering sparse regions.

1. **Acquisition of .hic files** Whenever available, hic files aligned to the hg38 reference genome were directly downloaded from the source database. If only hg19-aligned .hic files were provided, we converted them to hg38 using HiCLift (46). In cases where no .hic files existed, we generated them from raw fastq read files as described in the Supplementary Methods.
2. **TAD boundary calling** We identified TAD boundaries using OnTAD (47) at a 50 kb resolution with relaxed parameters (penalty=0, ldiff=0.25, lsize=2, minsz=3 and - maxsz=80) to ensure broad genome coverage for downstream structure modeling. Level 1 (outermost) TADs were selected as modeling units as they generally encompass complete domain structures.
3. **TAD-Based Segmentation of Modeling Regions** Modeling regions were defined by outermost TAD boundaries. Small TADs (<1Mb) were merged with neighboring domains if the combined size did not exceed 3.5 Mb, avoiding regions that are too small for reliable modeling.
4. **Filtering sparse regions** Regions with extremely low contact frequency, defined as < 1% of bins containing any contacts, were excluded. All contacts within such regions were removed from further analysis to improve modeling accuracy.

### Single-cell 3D Structure Reconstruction

We used CHROMATIX (48) to identify statistically significant folding reconstitutive (FoldRec) Hi-C interactions at 5 kb resolution. These interactions were used both for structural modeling and for visualization, enabling comparison with experimental Hi-C data (of similar resolution) and supporting downstream functional analysis. To generate sample-specific background models, we applied a fractal Monte Carlo approach to simulate large ensembles of chromatin fibers (500,000 conformations per sample) confined within the nuclear volume. These null ensembles incorporated only polymer physics and nuclear volume exclusion constraints, with nuclear sizes listed for each sample in Supplementary Table (49–74). Using a Bag of Little Bootstraps resampling approach, we derived null distributions of random chromatin contacts. Experimentally measured population Hi-C contacts were then compared to their corresponding null distributions, and interactions with a *p*.*adj* < 0.05 were retained as statistically significant FoldRec contacts. Single-cell chromatin structures were reconstructed from these FoldRec interactions using sBIF (39), which applies sequential Bayesian inference. For each dataset, we generated an ensemble of 5000 single-cell chromatin conformations. Within each model, two beads were considered to be in contact if their Euclidean distance was less than 80 nm (interpreted as a spherical distance threshold), consistent with prior studies (23, 39).

### Database Architecture and Web Implementation

ChromPolyerDB is implemented using a modern architecture. The backend is built with Flask, which provides RESTful APIs and handles core business logic. The frontend, developed in React with the Ant Design component library, delivers a consistent and responsive user interface. Data visualization is powered by D3.js for interactive 2D charts, an embedded IGV.js (75) genome browser provides track-based views of sequencing signals and annotations directly in the app, enabling intuitive exploration of biological datasets, and by Three.js for an in-browser 3D chromosome viewer that supports real-time interaction with structural models. All primary data is stored in a PostgreSQL database, enabling complex relational queries and scalability to large datasets. To improve performance, Redis is used as an in-memory caching layer, minimizing latency for frequently accessed data.

## RESULTS

### Database overview

ChromPolymerDB is a comprehensive, publicly accessible resource that hosts large-scale, high-resolution, single-cell 3D chromatin data, coupled with an interactive web interface for structural analysis and multi-omics integration capabilities (Figure 1). The database contains ∼10^8^ individual chromatin conformations reconstructed using sBIF, spanning more than 60,000 genomic regions at 5 kb resolution across 50 human cell types and experimental conditions (Supplementary Table S1). These datasets encompass 36 normal, healthy cell types and 14 disease samples, collectively representing 10 of the 11 major human physiological systems (excluding the urinary system). Beyond data access, ChromPolymerDB offers a suite of analytical tools, including locus-specific 3D visualization, structural measurements, and multi-omics integration, enabling both detailed single-sample interrogations and comparative analysis across cell types and conditions.

**Figure 1.**
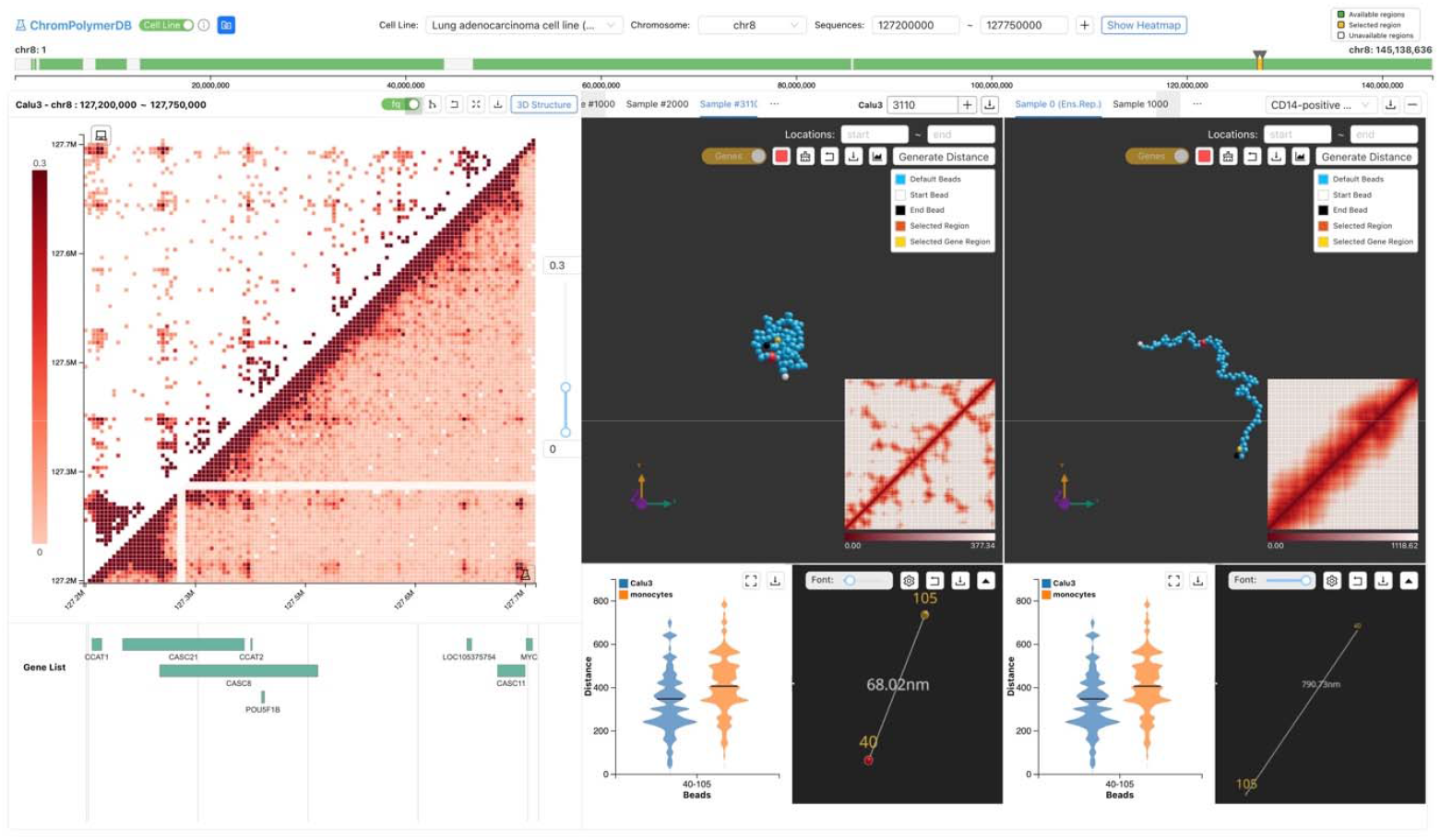
Overview of the ChromPolymerDB database. The example illustrates FoldRec-identified interactions and experimental Hi-C data in Calu3 cells, representative single-cell chromatin conformations from Calu3 and IMR90 cells, and calculated distance distributions between genomic elements of interest.

### Single-Sample Insights: Chromatin Architectures at the *MYC* Locus in Human Lung Cancer Cells

ChromPolymerDB enables detailed analysis of 3D chromatin organization within individual samples, supporting spatial regulatory investigations. Users can query statistically significant 2D folding interactions (FoldRec, see Methods) and reconstruct complete 3D genome structures.

As a case study, we analyzed the *MYC* locus (chr8: 127,200,000 – 127,750,000 bp) in Calu3 lung adenocarcinoma cells, which exhibit high *MYC* expression. *MYC*, a well-known oncogene, is regulated by multiple upstream enhancers (76–78) and plays a central role in tumor initiation and progression (79–81), making it an ideal target for chromatin architecture studies. Before reconstructing of single-cell 3D chromatin structures, we examined 2D Hi-C contact maps and epigenomic profiles to identify putative functional cis-regulatory elements. Users can define regions of interest by genomic coordinates or gene name (Figure 2A). ChromPolymerDB then displays the experiment-derived Hi-C heatmap of that locus (lower triangle) and CHROMATIX-derived FoldRec interactions (upper triangle) (Figure 2B). In Calu3 cells, the FoldRec interactions reveal contacts between the *MYC* promoter with multiple upstream regions, highlighting the potential regulatory significance of the chromatin organization in this region. Users can further overlay epigenomic tracks directly from the ENCODE Portal or upload their own custom tracks (e.g. histone modifications, transcription factor binding, or chromatin accessibility) for integrated analysis. This analysis enables identification of potential genomic loci of interest, such as regulatory elements, promoters, transcription factor binding sites or other functional regions, thereby facilitating an investigation of how these epigenetic features are related to the chromatin structure. For the *MYC* locus, DNase-seq, H3K27ac, H3K4me1, H3K4me3 and H3K27me3 ChIP-seq profiles, together with ChromHMM annotations, identified six putative regulatory enhancers (Figure 2D). Several have been experimentally linked to *MYC* regulation in cancers (78, 82), including prostate cancer (83, 84), B-cell malignancies (82), and colorectal cancer (85). RNA-seq data confirmed *MYC* overexpression in Calu3 cells (Figure 2C).

**Figure 2.**
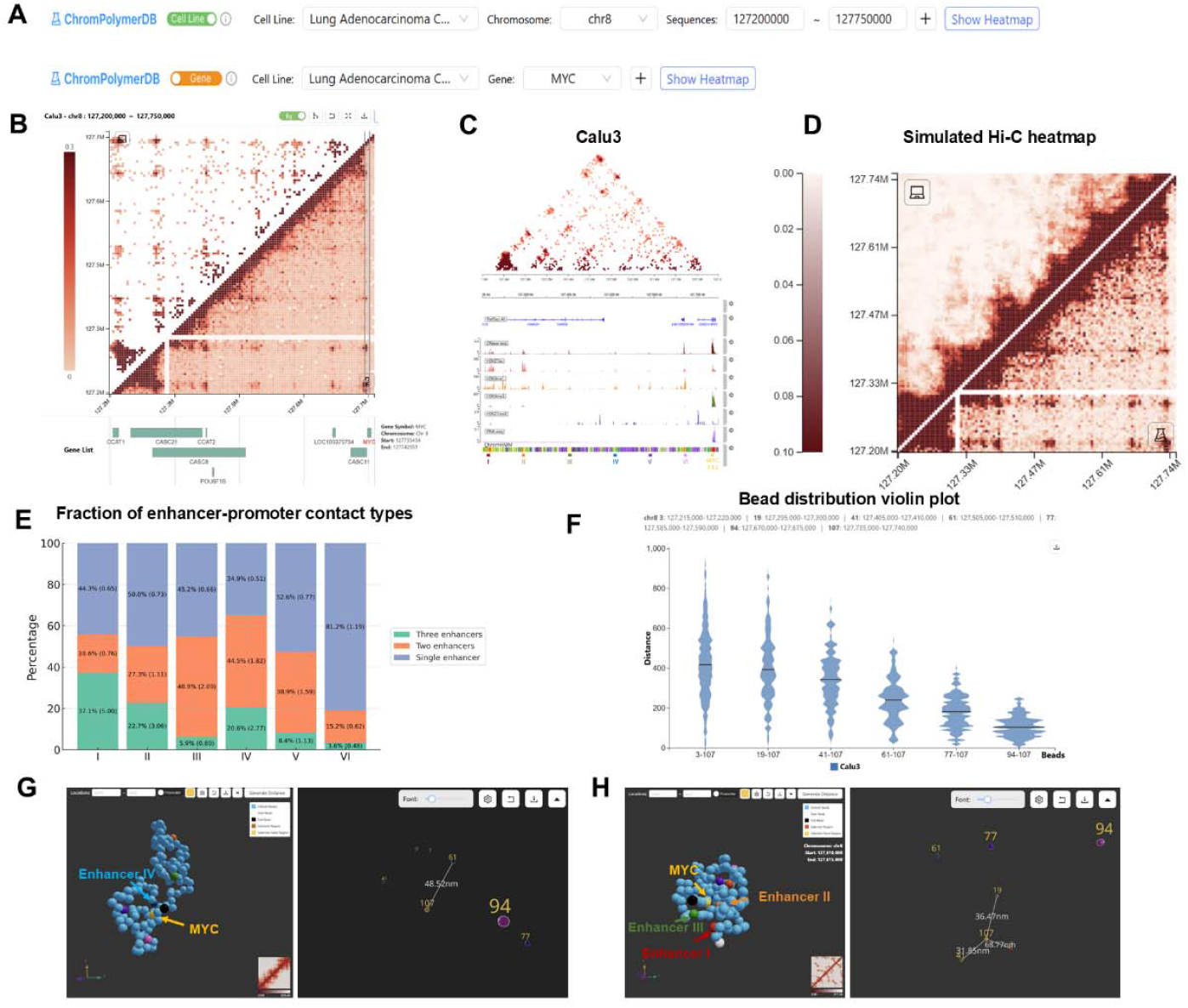
3D chromatin architecture at the *MYC* locus in Calu3 cells. (A) Two strategies for querying genomic regions (B) Hi-C heatmap of the *MYC* locus in Calu3 cells (lower triangle: experiment Hi-C; upper triangle: the FoldRec interactions). (C) FoldRec interactions overlaid with epigenomic profiles. Six putative enhancers (Enhancers I to VI) are underlined in red (chr8: 127,215,000-127,220,000), orange (chr8: 127,295,000-127,300,000), green (chr8: 127,400,000-127,405,000), blue (chr8: 127,505,000-127,510,000), purple (chr8: 127,585,000-127,590,000) and pink (chr8: 127,670,000-127,675,000). TSS is underlined in yellow (chr8: 127,735,000-127,740,000). (D) Comparison between simulated Hi-C heatmap (upper triangle) and experiment Hi-C heatmap (lower triangle). (E) Fraction of enhancer-promoter contact types for each putative enhancer, annotated with fold enrichment values. (F) Distribution of 3D distances between all putative enhancers and the *MYC* TSS across all generated structures. (G) Example single-cell chromatin structure with enhancer IV and the *MYC* TSS highlighted, illustrating close spatial proximity. (H) Example single-cell chromatin structure with three putative enhancers (red, orange, and green) and the *MYC* TSS (yellow) highlighted.

Using sBIF, we reconstructed 5,000 single-cell chromatin structures for this region, and the most representative single-cell conformations are displayed by default. Users can evaluate the accuracy of the reconstructed models by comparing the aggregate contact map derived from the simulated single-cell structures with the experimental Hi-C contact map (Figure 2D). The database enables users to switch seamlessly among all available structures for visualization or analysis. Distance measurements can be performed for any selected pair or group loci, such as enhancer-promoter pairs or multi-body contacts. For each selection the 3D distance in the displayed structure and the distribution of pairwise distances across the full set of 5,000 models can be calculated. To demonstrate the databases capabilities, we reconstructed single cell chromatin structures of the *MYC* locus (defined as described above) in the Calu3 cells. While bulk Hi-C maps suggest that all putative enhancers can contact the *MYC* TSS, our single-cell reconstructions reveal substantial heterogeneity in these interactions. To quantify this variability, we performed off-line analysis of the bead locations of the 5,000 reconstructed single-cell structures and found that 45.5% (2,275 in 5,000) exhibit at least one enhancer in contact with the *MYC* TSS. Of these 37.5% (1,871 in 5,000) structures had a single enhancer-promoter contact, 14.8% (336) had two enhancers in proximity, and 3% (68 in 5,000) show three enhancers contacting the *MYC* TSS. Thus, ∼18% of enhancer–promoter interactions occur in multibody configurations at the single-cell level, indicating that cooperative enhancer activity is a common feature of *MYC* regulation in these cells.

We further evaluated each putative enhancer’s tendency to participate in multibody interactions with the *MYC* TSS (Figure 2E). Enhancers I, III, and IV preferentially formed multibody contacts, with Enhancer I strongly enriched for three-enhancer interactions (Fold Enrichment = 5) and Enhancer III and IV favoring two-enhancer contacts (Fold Enrichment = 2 and 1.82). In contrast, Enhancer VI was more often involved in single enhancer–promoter interactions (Fold Enrichment = 1.19). Notably, most three-enhancer multibody interactions were formed by specific enhancer combinations such as (IV, V, VI) (54.4%), (I, IV, VI) (20.6%) and (I, II and III) (8.8%), suggesting that both long-range chromatin folding and local transcription factor binding may underlie cooperative *MYC* regulation.

To illustrate this variability, we highlight two representative structures. In one case (Figure 2G), only Enhancer IV is in close spatial proximity to the *MYC* promoter, with the remaining five enhancers located further away. In the second example (Figure 2H), three enhancers (I, II and III) cluster together with the *MYC* TSS, exemplifying a multibody configuration.

To further explore this spatial heterogeneity, users can perform ensemble-level clustering of single-cell structures. For the *MYC* locus, k-means clustering identified five distinct chromatin subgroups (Figure 3A; Supplementary Methods), each displaying unique structural patterns (Figure 3A, B). We also compared features such as the pairwise distance between Enhancer III and the *MYC* promoter (Figure 3C), as well as the radius of gyration of this region (Figure 3D), revealing substantial variation across the five subgroups. These structural differences may correspond to distinct regulatory states and could be integrated with other single-cell datasets, such as scRNA-seq or chromatin tracing (see, for example, Figure S1).

**Figure 3.**
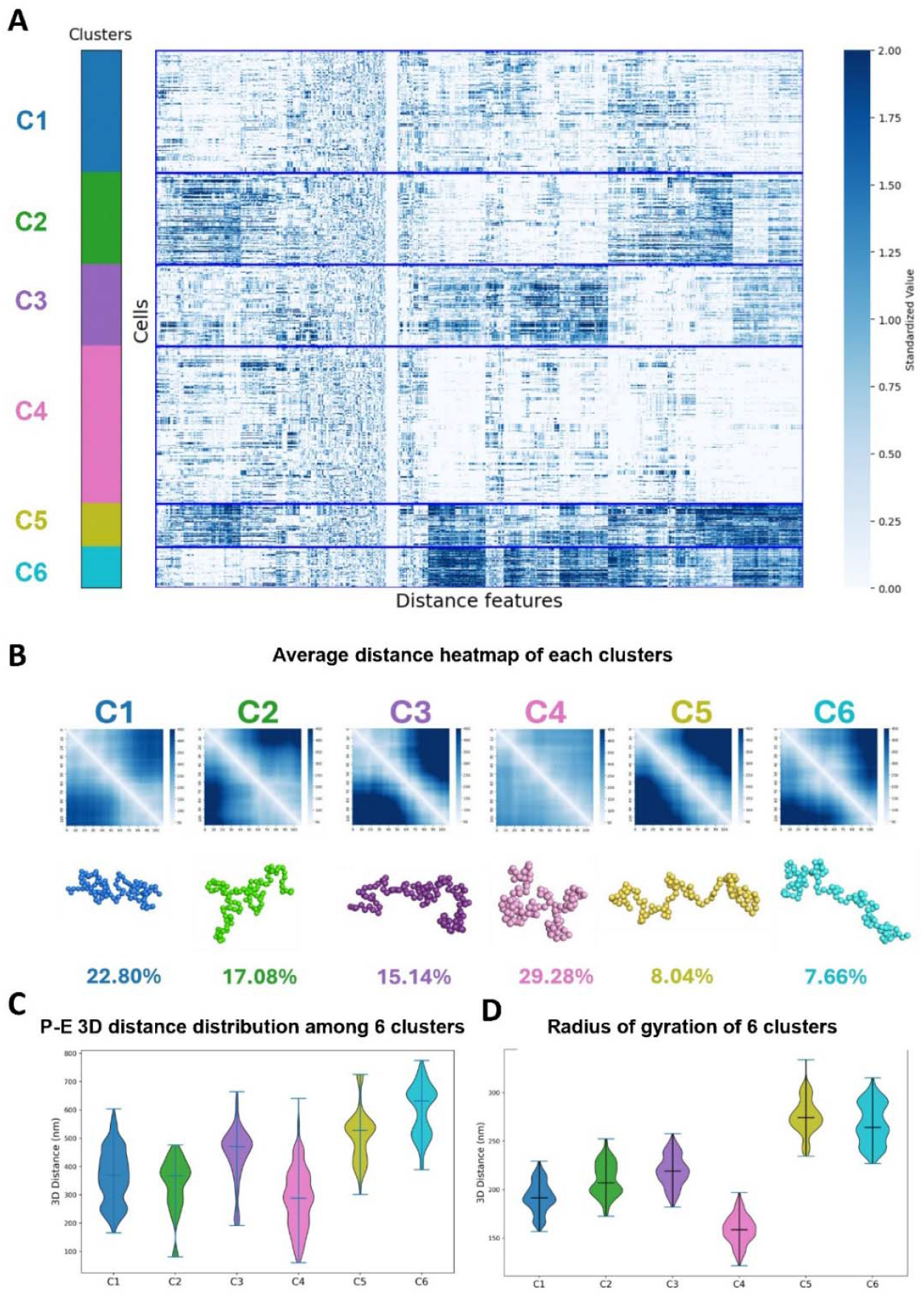
K-means clustering of single-cell chromatin structures of the *MYC* locus in Calu3 cells. (A) Heatmap of pairwise 3D distance features for all single-cell conformation of the *MYC* locus (B) Average 3D distance heatmaps for each of the five identified subclusters. (C) Distribution of 3D distances between putative enhancer III and the *MYC* TSS across the five subclusters. (D) Distribution of the radius of gyration for the *MYC* locus in each subcluster.

### Cross-Sample Comparison: Distinct Regulatory Mechanisms at the *MYC* Locus

ChromPolymerDB also enables direct comparison of chromatin structures across cell types, developmental stages, or disease states, providing a powerful framework for uncovering differences in regulatory mechanisms. The same structural analysis tools described in *Single-Sample Insights* can be applied to multi-sample datasets, allowing side-by-side evaluation of experimental Hi-C heatmaps, FoldRec interactions, and reconstructed 3D chromatin conformations.

As a demonstration, we compared *MYC* locus regulation in three cell types with contrasting *MYC* expression and epigenetic landscapes: Calu3 lung adenocarcinoma cells (aberrantly high *MYC* expression), GM12878 B-lymphoblastoid cells (15, 42) (high *MYC* expression), and primary CD14^+^ monocytes (43, 44) (negligible *MYC* expression). The observed epigenetic patterns are consistent with RNA-seq profiles displayed in Figure 4A. Epigenetic analysis (Figure 4A) revealed that Calu3 and GM12878 share two upstream enhancers (I and II), while Calu3 have four additional enhancers active (III, IV, V and VI), which are absent in the other two cell types. GM12878 harbors one unique enhancer (Enhancer VII), and monocytes contain two distinct enhancers that lack contact with the *MYC* TSS in the Hi-C data, suggesting that a regulatory relationship is unlikely, based on the available Hi-C data.

**Figure 4.**
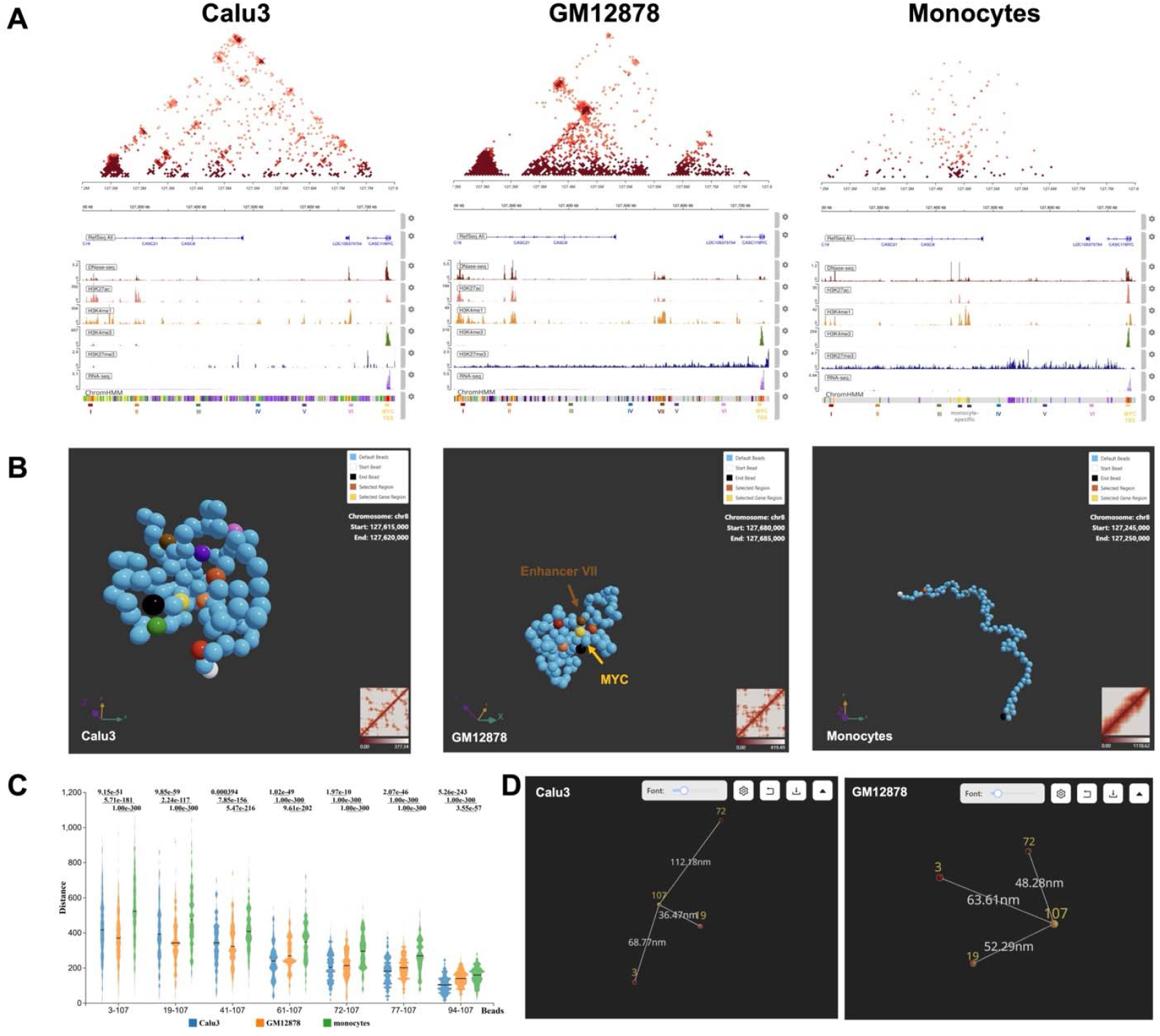
Case study of distinct regulatory mechanisms at the *MYC* Locus across cell types. (A) FoldRec interactions overlaid with epigenomic profiles for the *MYC* locus of Calu3 (left), GM12878 (middle) and monocytes (right). In Calu3 cells, six putative enhancers are underlined in red, orange, green, blue, purple and pink; the *MYC* TSS is underlined in yellow; and the GM12878-specific enhancer underlined in brown (chr8: 127,560,000-127,565,000). (B) Representative single-cell chromatin structures for Calu3 (left), GM12878 (middle) and monocytes (right). (C) Distributions of 3D distances between all seven putative enhancers and the *MYC* TSS across all generated structures in Calu3 (left), GM12878 (middle) and monocytes (right). (D) Comparison of 3D distances between GM12878 enhancers and the *MYC* TSS in Calu3 (left) and GM12878 (right) structures.

We reconstructed 5000 single-cell chromatin structures for all three cell types using sBIF to assess their spatial organization and multibody enhancer-promoter interaction patterns (Figure 4B-C). In both Calu3 and GM12878, active enhancers were positioned closer to the *MYC* TSS than in monocytes, consistent with the absence of active epigenetic marks and low *MYC* expression in the latter. However, the overall regulatory architectures differed markedly between the three cell types. (Figure 4A).

In Calu3, the most frequent multibody enhancer combinations were enriched, whereas these same configurations were 9.6-fold less frequent in GM12878, likely due to missing enhancer activities in that cell type. GM12878’s active enhancers (I, II and VII) occasionally colocalized with the *MYC* TSS in the single cell assemblies, but did not form stable multibody interactions, indicating a distinct mode of cooperative regulation to Calu3 cells (Figure 4D). Enhancer - Promoter contacts occurred in only ∼10% of GM12878 single-cell structures (446/5,000), compared to substantially higher frequencies in Calu3 cells. Among GM12878 contacts, 1.8% involved three enhancer and 10.3% involved two, underscoring the reduced prevalence of multibody configurations in GM12878 than Calu3.

These findings highlight ChromPolymerDB’s capacity to reveal cell type-specific regulatory mechanisms, demonstrating that even with similar *MYC* expression levels, Calu3 and GM12878 employ fundamentally different enhancer repertoires, degrees of multi-assembly, and cell-to-cell variability in chromatin organization.

## DISCUSSION

Recent advances in single-cell technologies have enabled high-resolution investigation of diverse biological modalities including transcription (86, 87), epigenetics modifications (88, 89), and chromatin structures (21, 27, 33). These approaches have transformed our understanding of cellular heterogeneity, revealing subpopulations and regulatory cell states that remain invisible in bulk analyses. However, generating accurate, high-resolution chromatin structures for individual single cells remains experimentally challenging and computationally intensive, limiting their accessibility to the broader research community.

To address these limitations, we developed sBIF to infer single-cell chromatin architectures from population Hi-C data (39). While sBIF achieves high structural resolution, its computational cost and technical complexity have restricted widespread adoption. ChromPolymerDB was therefore created as a publicly accessible, large-scale resource that delivers, high-resolution single-cell 3D chromatin structures alongside interactive tools for structural visualisation, analysis, and multi-omics integration.

Over the past years, several chromatin structure databases have been developed, greatly facilitating research in chromatin organization. For example, databases such as HiChIPdb (90), the 3D Genome Browser (91), 3DIV (92), and LoopCatalog (93) provide cross-species, high-resolution two-dimensional chromatin contact information at the bulk level, offering valuable support and laying the foundation for subsequent advances in chromatin structure research. However, the focus of these databases remains largely on two-dimensional bulk data, with limited coverage of three-dimensional structures and capacity to capture single-cell heterogeneity. In addition, databases such as GSDB (94), Nucleome Browser (95), and 3Disease Browser (96) have provided three-dimensional chromatin structure data with two-dimensional chromatin contact information, representing important progress toward more comprehensive and in-depth insights into chromatin organization. While highly valuable, these resources still have room for improvement in terms of resolution and sample coverage.

In contrast, ChromPolymerDB contains ∼10^8^ single-cell chromatin structures spanning more than 60,000 genomic regions at 5 Kb resolution, across 50 diverse human cell types and experimental conditions. This scale enables both in-depth analyses within a single cell type and systematic comparisons across multiple cellular contexts.

Utility of our database is illustrated through two case studies at the *MYC* locus. In human lung cancer Calu3 cells, single-cell reconstructions revealed extensive heterogeneity in enhancer-promoter spatial interactions that was obscured in bulk Hi-C data. This heterogeneity likely reflects regulatory variability relevant to *MYC* overexpression and provides hypothesis for targeted functional testing.

Beyond single sample analysis, ChromPolymerDB enables cross-sample comparisons to identify cell type-specific differences in regulatory architecture. Comparing *MYC* locus regulation in Calu3, GM12878, and primary CD14^+^ monocytes revealed marked contrast in enhancer usage, multibody interaction frequency, and spatial variability. These newly discovered differences could not be fully resolved from bulk Hi-C or epigenetic profiles alone. These findings underscore the critical role of single-cell structural data in uncovering the regulatory logic underlying gene expression.

By integrating such structural information with other genomic and epigenomic datasets, ChromPolymerDB will serve as a valuable platform that supports hypothesis generation, comparative 3D genomics, and mechanistic studies of gene regulation. Looking forward, we plan to expand the database to include additional species, incorporate new high-quality Hi-C datasets, and implement advanced structure analysis algorithms and more intuitive user tools. Integration with other single-cell omics modalities, (e.g., scRNA-seq and single-cell epigenomics) will further enable multi-modal analyses, offering a more comprehensive view of genome organization and regulation.

We anticipate that ChromPolymerDB will serve as a key community resource that can bridge the gap between raw chromatin conformation data and biological insight, thereby accelerating research in genome biology, regulatory genomics, and disease-associated chromatin remodeling.

## DATA AVAILABILITY

All the data is available from the URL: https://chrompolymerdb.bme.uic.edu/.

## SUPPLEMENTARY DATA

Supplementary Data are available at NAR online.

## AUTHOR CONTRIBUTIONS

Min Chen: Conceptualization, Investigation, Formal Analysis, Data Curation, Writing - Original Draft Preparation, Writing - Review & Editing. Lin Du: Conceptualization, Project Administration, Investigation, Formal Analysis, Writing -Original Draft Preparation, Writing - Review & Editing. Siyuan Zhao: Conceptualization, Investigation, Software, Data Curation Writing - Original Draft Preparation. Bowei Ye: Investigation. Pourya Delafrouz: Investigation. Hammad Farooq: Investigation. Debaleena ChattopadhyayL: Software. G. Elisabeta Marai: Software. Zhifeng Shao: Conceptualization, Writing - Review & Editing. Jie Liang: Conceptualization, Supervision, Funding Acquisition. Daniel M. Czajkowsky: Conceptualization, Writing - Review & Editing. Constantinos Chronis: Conceptualization, Supervision, Funding Acquisition, Writing - Review & Editing.

## ACKNOWLEDGEMENTS

We thank Aashik Mathew Prosper and Lavanya Vaddavalli for organizing the online tutorial section and Ming Cheng for suggestions.

## FUNDING

This work was supported by grants from the National Key R&D Program of China (No. 2020YFA0908100), the National Natural Science Foundation of China (Nos. 31971151, 81627801, 81972909, and 32370572), and the K.C. Wong Education Foundation (H.K.). Jie Liang was supported by National Institutes of Health R03OD036492 and R35GM127084. Constantinos Chronis was supported by National Institutes of Health P01 HL160469 and National Institutes of Health R01HL170286 grants. This work was also supported by INCITE Program awards to Jie Liang and Constantinos Chronis. Siyuan Zhao and G. Elisabeta Marai were supported by National Institutes of Health National Institutes of Health NCI R01CA258827, National Institutes of Health UG3 TR004501, National Science Foundation CNS-2320261, and the UIC Institute for Health Data Science Research.

## CONFLICT OF INTEREST

The authors have declared no competing interests.

## References

1. Venkatesh, S. and Workman, J.L. (2015) Histone exchange, chromatin structure and the regulation of transcription. Nat Rev Mol Cell Biol, 16, 178–189.

2. Sexton, T. and Cavalli, G. (2015) The Role of Chromosome Domains in Shaping the Functional Genome. Cell, 160, 1049–1059.

3. Rowley, M.J. and Corces, V.G. (2018) Organizational principles of 3D genome architecture. Nat Rev Genet, 19, 789–800.

4. Dinant, C., Houtsmuller, A.B. and Vermeulen, W. (2008) Chromatin structure and DNA damage repair. Epigenetics & Chromatin, 1, 9.

5. Dixon, J.R., Jung, I., Selvaraj, S., Shen, Y., Antosiewicz-Bourget, J.E., Lee, A.Y., Ye, Z., Kim, A., Rajagopal, N., Xie, W., et al. (2015) Chromatin architecture reorganization during stem cell differentiation. Nature, 518, 331–336.

6. Zhang, G., Li, Y. and Wei, G. (2023) Multi-omic analysis reveals dynamic changes of three-dimensional chromatin architecture during T cell differentiation. Commun Biol, 6, 773.

7. Zheng, H. and Xie, W. (2019) The role of 3D genome organization in development and cell differentiation. Nat Rev Mol Cell Biol, 20, 535–550.

8. Bonev, B., Mendelson Cohen, N., Szabo, Q., Fritsch, L., Papadopoulos, G.L., Lubling, Y., Xu, X., Lv, X., Hugnot, J.-P., Tanay, A., et al. (2017) Multiscale 3D Genome Rewiring during Mouse Neural Development. Cell, 171, 557-572.e24.

9. Hug, C.B., Grimaldi, A.G., Kruse, K. and Vaquerizas, J.M. (2017) Chromatin Architecture Emerges during Zygotic Genome Activation Independent of Transcription. Cell, 169, 216-228.e19.

10. Xu, J., Song, F., Lyu, H., Kobayashi, M., Zhang, B., Zhao, Z., Hou, Y., Wang, X., Luan, Y., Jia, B., et al. (2022) Subtype-specific 3D genome alteration in acute myeloid leukaemia. Nature, 611, 387–398.

11. Xu, Z., Lee, D.-S., Chandran, S., Le, V.T., Bump, R., Yasis, J., Dallarda, S., Marcotte, S., Clock, B., Haghani, N., et al. (2022) Structural variants drive context-dependent oncogene activation in cancer. Nature, 612, 564–572.

12. Franke, M., Ibrahim, D.M., Andrey, G., Schwarzer, W., Heinrich, V., Schöpflin, R., Kraft, K., Kempfer, R., Jerković, I., Chan, W.-L., et al. (2016) Formation of new chromatin domains determines pathogenicity of genomic duplications. Nature, 538, 265–269.

13. Lupiáñez, D.G., Kraft, K., Heinrich, V., Krawitz, P., Brancati, F., Klopocki, E., Horn, D., Kayserili, H., Opitz, J.M., Laxova, R., et al. (2015) Disruptions of Topological Chromatin Domains Cause Pathogenic Rewiring of Gene-Enhancer Interactions. Cell, 161, 1012–1025.

14. Perez-Rathke, A., Mali, S., Du, L. and Liang, J. (2019) Alterations in Chromatin Folding Patterns in Cancer Variant-Enriched Loci. In 2019 IEEE EMBS International Conference on Biomedical & Health Informatics (BHI). IEEE, Chicago, IL, USA, pp. 1–4.

15. Rao, S.S.P., Huntley, M.H., Durand, N.C., Stamenova, E.K., Bochkov, I.D., Robinson, J.T., Sanborn, A.L., Machol, I., Omer, A.D., Lander, E.S., et al. (2014) A 3D map of the human genome at kilobase resolution reveals principles of chromatin looping. Cell, 159, 1665–1680.

16. Lieberman-Aiden, E., Van Berkum, N.L., Williams, L., Imakaev, M., Ragoczy, T., Telling, A., Amit, I., Lajoie, B.R., Sabo, P.J., Dorschner, M.O., et al. (2009) Comprehensive Mapping of Long-Range Interactions Reveals Folding Principles of the Human Genome. Science, 326, 289–293.

17. Hsieh, T.-H.S., Weiner, A., Lajoie, B., Dekker, J., Friedman, N. and Rando, O.J. (2015) Mapping Nucleosome Resolution Chromosome Folding in Yeast by Micro-C. Cell, 162, 108–119.

18. Beagrie, R.A., Scialdone, A., Schueler, M., Kraemer, D.C.A., Chotalia, M., Xie, S.Q., Barbieri, M., De Santiago, I., Lavitas, L.-M., Branco, M.R., et al. (2017) Complex multi-enhancer contacts captured by genome architecture mapping. Nature, 543, 519–524.

19. Quinodoz, S.A., Ollikainen, N., Tabak, B., Palla, A., Schmidt, J.M., Detmar, E., Lai, M.M., Shishkin, A.A., Bhat, P., Takei, Y., et al. (2018) Higher-Order Inter-chromosomal Hubs Shape 3D Genome Organization in the Nucleus. Cell, 174, 744-757.e24.

20. Cremer, T. and Cremer, C. (2001) Chromosome territories, nuclear architecture and gene regulation in mammalian cells. Nat Rev Genet, 2, 292–301.

21. Bintu, B., Mateo, L.J., Su, J.-H., Sinnott-Armstrong, N.A., Parker, M., Kinrot, S., Yamaya, K., Boettiger, A.N. and Zhuang, X. (2018) Super-resolution chromatin tracing reveals domains and cooperative interactions in single cells. Science, 362, eaau1783.

22. Wang, Q., Sun, Q., Czajkowsky, D.M. and Shao, Z. (2018) Sub-kb Hi-C in D. melanogaster reveals conserved characteristics of TADs between insect and mammalian cells. Nat Commun, 9, 188.

23. Du, L., Farooq, H., Delafrouz, P. and Liang, J. (2025) Structural basis of differential gene expression at eQTLs loci from high-resolution ensemble models of 3D single-cell chromatin conformations. Bioinformatics, 41, btaf050.

24. Liu, X., Sun, Q., Wang, Q., Hu, C., Chen, X., Li, H., Czajkowsky, D.M. and Shao, Z. (2022) Epithelial Cells in 2D and 3D Cultures Exhibit Large Differences in Higher-Order Genomic Interactions. Genomics, Proteomics & Bioinformatics, 20, 101–109.

25. Liu, Y., Li, H., Czajkowsky, D.M. and Shao, Z. (2021) Monocytic THP-1 cells diverge significantly from their primary counterparts: a comparative examination of the chromosomal conformations and transcriptomes. Hereditas, 158, 43.

26. Liu, Y., Chen, M., Liu, X., Xu, Z., Li, X., Guo, Y., Czajkowsky, D.M. and Shao, Z. (2025) Histo-LCM-Hi-C reveals the 3D chromatin conformation of spatially localized rare cells in tissues at high resolution. 10.1101/2025.06.10.658807.

27. Nagano, T., Lubling, Y., Stevens, T.J., Schoenfelder, S., Yaffe, E., Dean, W., Laue, E.D., Tanay, A. and Fraser, P. (2013) Single-cell Hi-C reveals cell-to-cell variability in chromosome structure. Nature, 502, 59–64.

28. Nagano, T., Lubling, Y., Várnai, C., Dudley, C., Leung, W., Baran, Y., Mendelson Cohen, N., Wingett, S., Fraser, P. and Tanay, A. (2017) Cell-cycle dynamics of chromosomal organization at single-cell resolution. Nature, 547, 61–67.

29. Shi, G. and Thirumalai, D. (2019) Conformational heterogeneity in human interphase chromosome organization reconciles the FISH and Hi-C paradox. Nat Commun, 10, 3894.

30. Liang, J. and Perez-Rathke, A. (2021) Minimalistic 3D chromatin models: Sparse interactions in single cells drive the chromatin fold and form many-body units. Current Opinion in Structural Biology, 71, 200–214.

31. Finn, E.H., Pegoraro, G., Brandão, H.B., Valton, A.-L., Oomen, M.E., Dekker, J., Mirny, L. and Misteli, T. (2019) Extensive Heterogeneity and Intrinsic Variation in Spatial Genome Organization. Cell, 176, 1502-1515.e10.

32. Ramani, V., Deng, X., Qiu, R., Gunderson, K.L., Steemers, F.J., Disteche, C.M., Noble, W.S., Duan, Z. and Shendure, J. (2017) Massively multiplex single-cell Hi-C. Nat Methods, 14, 263–266.

33. Tan, L., Xing, D., Chang, C.-H., Li, H. and Xie, X.S. (2018) Three-dimensional genome structures of single diploid human cells. Science, 361, 924–928.

34. Flyamer, I.M., Gassler, J., Imakaev, M., Brandão, H.B., Ulianov, S.V., Abdennur, N., Razin, S.V., Mirny, L.A. and Tachibana-Konwalski, K. (2017) Single-nucleus Hi-C reveals unique chromatin reorganization at oocyte-to-zygote transition. Nature, 544, 110–114.

35. Li, X., Zeng, G., Li, A. and Zhang, Z. (2021) DeTOKI identifies and characterizes the dynamics of chromatin TAD-like domains in a single cell. Genome Biol, 22, 217.

36. Yu, M., Abnousi, A., Zhang, Y., Li, G., Lee, L., Chen, Z., Fang, R., Lagler, T.M., Yang, Y., Wen, J., et al. (2021) SnapHiC: a computational pipeline to identify chromatin loops from single-cell Hi-C data. Nat Methods, 18, 1056–1059.

37. Zhang, R., Zhou, T. and Ma, J. (2022) Ultrafast and interpretable single-cell 3D genome analysis with Fast-Higashi. Cell Systems, 13, 798-807.e6.

38. Zhou, J., Ma, J., Chen, Y., Cheng, C., Bao, B., Peng, J., Sejnowski, T.J., Dixon, J.R. and Ecker, J.R. (2019) Robust single-cell Hi-C clustering by convolution- and random-walk–based imputation. Proc. Natl. Acad. Sci. U.S.A., 116, 14011–14018.

39. Sun, Q., Perez-Rathke, A., Czajkowsky, D.M., Shao, Z. and Liang, J. (2021) High-resolution single-cell 3D-models of chromatin ensembles during Drosophila embryogenesis. Nat Commun, 12, 205.

40. Farooq, H., Du, L., Delafrouz, P., Jiang, W., Chronis, C. and Liang, J. (2024) A New Approach for Discovering Functional Links Connecting Non-Coding Regulatory Variants to Gene Targets. 10.1101/2024.06.13.598913.

41. Delafrouz, P., Farooq, H., Du, L., Ma, A. and Liang, J. (2025) Effects of Lamina-Chromatin Attachment on Super Long-Range Chromatin Interactions. 10.1101/2025.02.13.638183.

42. Reiff, S.B., Schroeder, A.J., Kırlı, K., Cosolo, A., Bakker, C., Mercado, L., Lee, S., Veit, A.D., Balashov, A.K., Vitzthum, C., et al. (2022) The 4D Nucleome Data Portal as a resource for searching and visualizing curated nucleomics data. Nat Commun, 13, 2365.

43. Jou, J., Gabdank, I., Luo, Y., Lin, K., Sud, P., Myers, Z., Hilton, J.A., Kagda, M.S., Lam, B., O’Neill, E., et al. (2019) The ENCODE Portal as an Epigenomics Resource. CP in Bioinformatics, 68, e89.

44. The ENCODE Project Consortium (2012) An integrated encyclopedia of DNA elements in the human genome. Nature, 489, 57–74.

45. Clough, E. and Barrett, T. (2016) The Gene Expression Omnibus Database. In Mathé, E., Davis, S. (eds), Statistical Genomics, Methods in Molecular Biology. Springer New York, New York, NY, Vol. 1418, pp. 93–110.

46. Wang, X. and Yue, F. (2023) HiCLift: a fast and efficient tool for converting chromatin interaction data between genome assemblies. Bioinformatics, 39, btad389.

47. An, L., Yang, T., Yang, J., Nuebler, J., Xiang, G., Hardison, R.C., Li, Q. and Zhang, Y. (2019) OnTAD: hierarchical domain structure reveals the divergence of activity among TADs and boundaries. Genome Biol, 20, 282.

48. Perez-Rathke, A., Sun, Q., Wang, B., Boeva, V., Shao, Z. and Liang, J. (2020) CHROMATIX: computing the functional landscape of many-body chromatin interactions in transcriptionally active loci from deconvolved single cells. Genome Biol, 21, 13.

49. Sanborn, A.L., Rao, S.S.P., Huang, S.-C., Durand, N.C., Huntley, M.H., Jewett, A.I., Bochkov, I.D., Chinnappan, D., Cutkosky, A., Li, J., et al. (2015) Chromatin extrusion explains key features of loop and domain formation in wild-type and engineered genomes. Proc Natl Acad Sci U S A, 112, E6456–6465.

50. Ibarra, J., Hershenhouse, T., Almassalha, L., Walterhouse, D., Backman, V. and MacQuarrie, K.L. (2023) Differentiation-dependent chromosomal organization changes in normal myogenic cells are absent in rhabdomyosarcoma cells. Front. Cell Dev. Biol., 11, 1293891.

51. Lawton, S., Danzman, R.A., Spagnuolo, R., Stephan, S., Graul, S., Wiggan, O., Ghosh, S. and Prasad, A. (2024) Cell Morphology accurately predicts the nuclear shape of adherent cells. 10.1101/2024.12.28.630588.

52. Meng, L. (2023) Chromatin-modifying enzymes as modulators of nuclear size during lineage differentiation. Cell Death Discov., 9, 384.

53. Bueno, C., García-Bernal, D., Martínez, S., Blanquer, M. and Moraleda, J.M. (2024) The nuclei of human adult stem cells can move within the cell and generate cellular protrusions to contact other cells. Stem Cell Res Ther, 15, 32.

54. Sadeghi Shoreh Deli, A., Scharf, S., Steiner, Y., Bein, J., Hansmann, M.-L. and Hartmann, S. (2022) 3D analyses reveal T cells with activated nuclear features in T-cell/histiocyte-rich large B-cell lymphoma. Modern Pathology, 35, 1431–1438.

55. Dorland, Y.L., Cornelissen, A.S., Kuijk, C., Tol, S., Hoogenboezem, M., Van Buul, J.D., Nolte, M.A., Voermans, C. and Huveneers, S. (2019) Nuclear shape, protrusive behaviour and in vivo retention of human bone marrow mesenchymal stromal cells is controlled by Lamin-A/C expression. Sci Rep, 9, 14401.

56. Phuangbubpha, P., Thara, S., Sriboonaied, P., Saetan, P., Tumnoi, W. and Charoenpanich, A. (2023) Optimizing THP-1 Macrophage Culture for an Immune-Responsive Human Intestinal Model. Cells, 12, 1427.

57. Lee, S.Y., Lloyd, W.R., Chandra, M., Wilson, R.H., McKenna, B., Simeone, D., Scheiman, J. and Mycek, M.-A. (2013) Characterizing human pancreatic cancer precursor using quantitative tissue optical spectroscopy. Biomed. Opt. Express, 4, 2828.

58. Rowat, A.C., Jaalouk, D.E., Zwerger, M., Ung, W.L., Eydelnant, I.A., Olins, D.E., Olins, A.L., Herrmann, H., Weitz, D.A. and Lammerding, J. (2013) Nuclear Envelope Composition Determines the Ability of Neutrophil-type Cells to Passage through Micron-scale Constrictions. Journal of Biological Chemistry, 288, 8610–8618.

59. Kuroda, M., Tanabe, H., Yoshida, K., Oikawa, K., Saito, A., Kiyuna, T., Mizusawa, H. and Mukai, K. (2004) Alteration of chromosome positioning during adipocyte differentiation. Journal of Cell Science, 117, 5897–5903.

60. Toft, M.H., Gredal, O. and Pakkenberg, B. (2005) The size distribution of neurons in the motor cortex in amyotrophic lateral sclerosis. Journal of Anatomy, 207, 399–407.

61. Khalo, I.V., Konokhova, A.I., Orlova, D.Y., Trusov, K.V., Yurkin, M.A., Bartova, E., Kozubek, S., Maltsev, V.P. and Chernyshev, A.V. (2018) Nuclear apoptotic volume decrease in individual cells: Confocal microscopy imaging and kinetic modeling. Journal of Theoretical Biology, 454, 60–69.

62. Maul, G. and Deaven, L. (1977) Quantitative determination of nuclear pore complexes in cycling cells with differing DNA content. The Journal of Cell Biology, 73, 748–760.

63. Aru, A. and Nielsen, K. (1989) Stereological Estimates of Nuclear Volume in Primary Lung Cancer. Pathology - Research and Practice, 185, 735–739.

64. Lim, J.Y., Hansen, J.C., Siedlecki, C.A., Runt, J. and Donahue, H.J. (2005) Human foetal osteoblastic cell response to polymer-demixed nanotopographic interfaces. J. R. Soc. Interface., 2, 97–108.

65. Mayhew, T.M., Pharaoh, A., Austin, A. and Fagan, D.G. (1997) Stereological estimates of nuclear number in human ventricular cardiomyocytes before and after birth obtained using physical disectors. Journal of Anatomy, 191, 107–115.

66. Qusous, A. and Kerrigan, M.J.P. (2012) Quantification of Changes in Morphology, Mechanotransduction, and Gene Expression in Bovine Articular Chondrocytes in Response to 2-Dimensional Culture Indicates the Existence of a Novel Phenotype. CARTILAGE, 3, 222–234.

67. Ganguly, A., Bhattacharjee, C., Bhave, M., Kailaje, V., Jain, B.K., Sengupta, I., Rangarajan, A. and Bhattacharyya, D. (2016) Perturbation of nucleo-cytoplasmic transport affects size of nucleus and nucleolus in human cells. FEBS Letters, 590, 631–643.

68. Tang, F., Du, Q. and Liu, Y.-J. (2010) Plasmacytoid dendritic cells in antiviral immunity and autoimmunity. Sci. China Life Sci., 53, 172–182.

69. Seaman, L., Meixner, W., Snyder, J. and Rajapakse, I. (2015) Periodicity of nuclear morphology in human fibroblasts. Nucleus, 6, 408–416.

70. Feng, Y., Zhang, N., Jacobs, K.M., Jiang, W., Yang, L.V., Li, Z., Zhang, J., Lu, J.Q. and Hu, X. (2014) Polarization imaging and classification of J urkat T and R amos B cells using a flow cytometer. Cytometry Pt A, 85, 817–826.

71. Hancock, R. and Hadj-Sahraoui, Y. (2009) Isolation of Cell Nuclei Using Inert Macromolecules to Mimic the Crowded Cytoplasm. PLoS ONE, 4, e7560.

72. Uyama, N., Uchihara, T., Mochizuki, Y., Nakamura, A., Takahashi, R. and Mizutani, T. (2009) Selective Nuclear Shrinkage of Oligodendrocytes Lacking Glial Cytoplasmic Inclusions in Multiple System Atrophy: A 3-Dimensional Volumetric Study. J Neuropathol Exp Neurol, 68, 1084–1091.

73. Hansen, E., Rolling, C., Wang, M. and Holaska, J.M. (2024) Emerin deficiency drives MCF7 cells to an invasive phenotype. 10.1101/2024.02.21.581379.

74. Deng, W., Chen, M., Tang, Y., Zhang, L., Xu, Z., Li, X., Czajkowsky, D.M. and Shao, Z. (2022) Temporal Analysis Reveals the Transient Differential Expression of Transcription Factors That Underlie the Trans-Differentiation of Human Monocytes to Macrophages. IJMS, 23, 15830.

75. Robinson, J.T., Thorvaldsdottir, H., Turner, D. and Mesirov, J.P. (2023) igv.js: an embeddable JavaScript implementation of the Integrative Genomics Viewer (IGV). Bioinformatics, 39, btac830.

76. MYC activates transcriptional enhancers to drive cancer progression (2024) Nat Genet, 56, 567–568.

77. Jia, Y., Chng, W.-J. and Zhou, J. (2019) Super-enhancers: critical roles and therapeutic targets in hematologic malignancies. J Hematol Oncol, 12, 77.

78. Lancho, O. and Herranz, D. (2018) The MYC Enhancer-ome: Long-Range Transcriptional Regulation of MYC in Cancer. Trends in Cancer, 4, 810–822.

79. Dhanasekaran, R., Deutzmann, A., Mahauad-Fernandez, W.D., Hansen, A.S., Gouw, A.M. and Felsher, D.W. (2022) The MYC oncogene — the grand orchestrator of cancer growth and immune evasion. Nat Rev Clin Oncol, 19, 23–36.

80. Kalkat, M., De Melo, J., Hickman, K., Lourenco, C., Redel, C., Resetca, D., Tamachi, A., Tu, W. and Penn, L. (2017) MYC Deregulation in Primary Human Cancers. Genes, 8, 151.

81. Gabay, M., Li, Y. and Felsher, D.W. (2014) MYC Activation Is a Hallmark of Cancer Initiation and Maintenance. Cold Spring Harbor Perspectives in Medicine, 4, a014241–a014241.

82. Hnisz, D., Abraham, B.J., Lee, T.I., Lau, A., Saint-André, V., Sigova, A.A., Hoke, H.A. and Young, R.A. (2013) Super-Enhancers in the Control of Cell Identity and Disease. Cell, 155, 934–947.

83. Wasserman, N.F., Aneas, I. and Nobrega, M.A. (2010) An 8q24 gene desert variant associated with prostate cancer risk confers differential in vivo activity to a MYC enhancer. Genome Res., 20, 1191–1197.

84. Jia, L., Landan, G., Pomerantz, M., Jaschek, R., Herman, P., Reich, D., Yan, C., Khalid, O., Kantoff, P., Oh, W., et al. (2009) Functional Enhancers at the Gene-Poor 8q24 Cancer-Linked Locus. PLoS Genet, 5, e1000597.

85. Pomerantz, M.M., Ahmadiyeh, N., Jia, L., Herman, P., Verzi, M.P., Doddapaneni, H., Beckwith, C.A., Chan, J.A., Hills, A., Davis, M., et al. (2009) The 8q24 cancer risk variant rs6983267 shows long-range interaction with MYC in colorectal cancer. Nat Genet, 41, 882–884.

86. Jovic, D., Liang, X., Zeng, H., Lin, L., Xu, F. and Luo, Y. (2022) Single-cell RNA sequencing technologies and applications: A brief overview. Clinical & Translational Med, 12, e694.

87. Cloonan, N., Forrest, A.R.R., Kolle, G., Gardiner, B.B.A., Faulkner, G.J., Brown, M.K., Taylor, D.F., Steptoe, A.L., Wani, S., Bethel, G., et al. (2008) Stem cell transcriptome profiling via massive-scale mRNA sequencing. Nat Methods, 5, 613–619.

88. Cusanovich, D.A., Hill, A.J., Aghamirzaie, D., Daza, R.M., Pliner, H.A., Berletch, J.B., Filippova, G.N., Huang, X., Christiansen, L., DeWitt, W.S., et al. (2018) A Single-Cell Atlas of In Vivo Mammalian Chromatin Accessibility. Cell, 174, 1309-1324.e18.

89. Kelsey, G., Stegle, O. and Reik, W. (2017) Single-cell epigenomics: Recording the past and predicting the future. Science, 358, 69–75.

90. Zeng, W., Liu, Q., Yin, Q., Jiang, R. and Wong, W.H. (2023) HiChIPdb: a comprehensive database of HiChIP regulatory interactions. Nucleic Acids Research, 51, D159–D166.

91. Wang, Y., Song, F., Zhang, B., Zhang, L., Xu, J., Kuang, D., Li, D., Choudhary, M.N.K., Li, Y., Hu, M., et al. (2018) The 3D Genome Browser: a web-based browser for visualizing 3D genome organization and long-range chromatin interactions. Genome Biol, 19, 151.

92. Yang, D., Jang, I., Choi, J., Kim, M.-S., Lee, A.J., Kim, H., Eom, J., Kim, D., Jung, I. and Lee, B. (2018) 3DIV: A 3D-genome Interaction Viewer and database. Nucleic Acids Research, 46, D52–D57.

93. Reyna, J., Fetter, K., Ignacio, R., Ali Marandi, C.C., Ma, A., Rao, N., Jiang, Z., Figueroa, D.S., Bhattacharyya, S. and Ay, F. (2024) Loop Catalog: a comprehensive HiChIP database of human and mouse samples. 10.1101/2024.04.26.591349.

94. Oluwadare, O., Highsmith, M., Turner, D., Lieberman Aiden, E. and Cheng, J. (2020) GSDB: a database of 3D chromosome and genome structures reconstructed from Hi-C data. BMC Mol and Cell Biol, 21, 60.

95. Zhu, X., Zhang, Y., Wang, Y., Tian, D., Belmont, A.S., Swedlow, J.R. and Ma, J. (2022) Nucleome Browser: an integrative and multimodal data navigation platform for 4D Nucleome. Nat Methods, 19, 911–913.

96. Li, R., Liu, Y., Li, T. and Li, C. (2016) 3Disease Browser: A Web server for integrating 3D genome and disease-associated chromosome rearrangement data. Sci Rep, 6, 34651.

