## Supplemental info for "ChromPolymerDB: A High-Resolution Database of Single-Cell 3D Chromatin Structures for Functional Genomics"

**Supplementary**

**Methods**

**Construction of .hic files from .fastq files**

For samples lacking pre-generated .hic files, we constructed them directly from raw .fastq read files. Hi-C sequencing reads were aligned to the hg38 reference genome using BWA (1) (0.7.18) with “-SP5M” option. Processed read pairs were generated using Pairtools (2), with removal of low-quality or invalid alignments. Specifically, PCR duplicates, multi-mapped reads, read pairs with mapping quality (MAPQ) below 30, and reads mapped to sex chromosomes, alternative contigs, or mitochondrial genome were removed. Additionally, Hi-C artifacts such as non-specific ligations (fragments with sizes above 600 bp), dangling ends (inward-facing read pairs less than 1 kb apart), self-ligations (outward-facing read pairs less than 25 kb apart), and mirror reads (reads mapped to the same strand of the same restriction fragment) were also removed. Only valid read pairs were retained for downstream processing. .hic files were generated using Juicer tools (v1.19.02) (3), applying Knight–Ruiz (KR) normalization by default. In cases where KR normalization failed, vanilla coverage (VC) normalization. Basically, all analysis were performed on KR-normalized data by default. In cases where KR normalization failed, VC normalization was used instead.

**Single-cell 3D Structure Clustering**

With downloaded structural data, pairwise distances between all bead pairs were first computed for each single-cell structure to generate inter-locus distance matrices. These matrices were then standardized via feature-wise Z-score normalization to center each distance measurement at zero mean and unit variance. The resulting standardized data were clustered into six discrete subpopulations using the K-means algorithm [k = 6, determined based on the elbow point of the inertia curve; implemented in Python(3.11.7) with scikit-learn(1.2.2) (4); random_state = 42]. To uncover relationships among distance features, Euclidean distances between columns were calculated and subjected to agglomerative hierarchical clustering with Ward’s minimum-variance linkage [performed in Python(3.11.7) using scikit-learn(1.2.2)], and the dendrogram order was used to permute the heatmap’s column arrangement.

**Radius of gyration of single-cell 3D chromatin structures**

For each single-cell chromatin conformation, we quantified structural compactness by computing the radius of gyration ($R_{g}$) directly from the inter-bead distance matrix, thereby avoiding explicit reconstruction of the center of mass. Given an N×N symmetric distance matrix D with elements $d_{ij}$ (and $d_{ii}$=0), we used the identity

$$R_{g}^{2}=\frac{1}{2N^{2}}\sum_{i=1}^{N} \sum_{j=1}^{N} d_{ij}^{2}$$

We implemented this calculation in Python by squaring all pairwise distances, summing over the matrix, normalizing by 2N^2^, and taking the square root to obtain$R_{g}$. Because the method depends only on pairwise separations, it is invariant to rigid translations and rotations of the configuration; units of $R_{g}$ match those of the input distances.

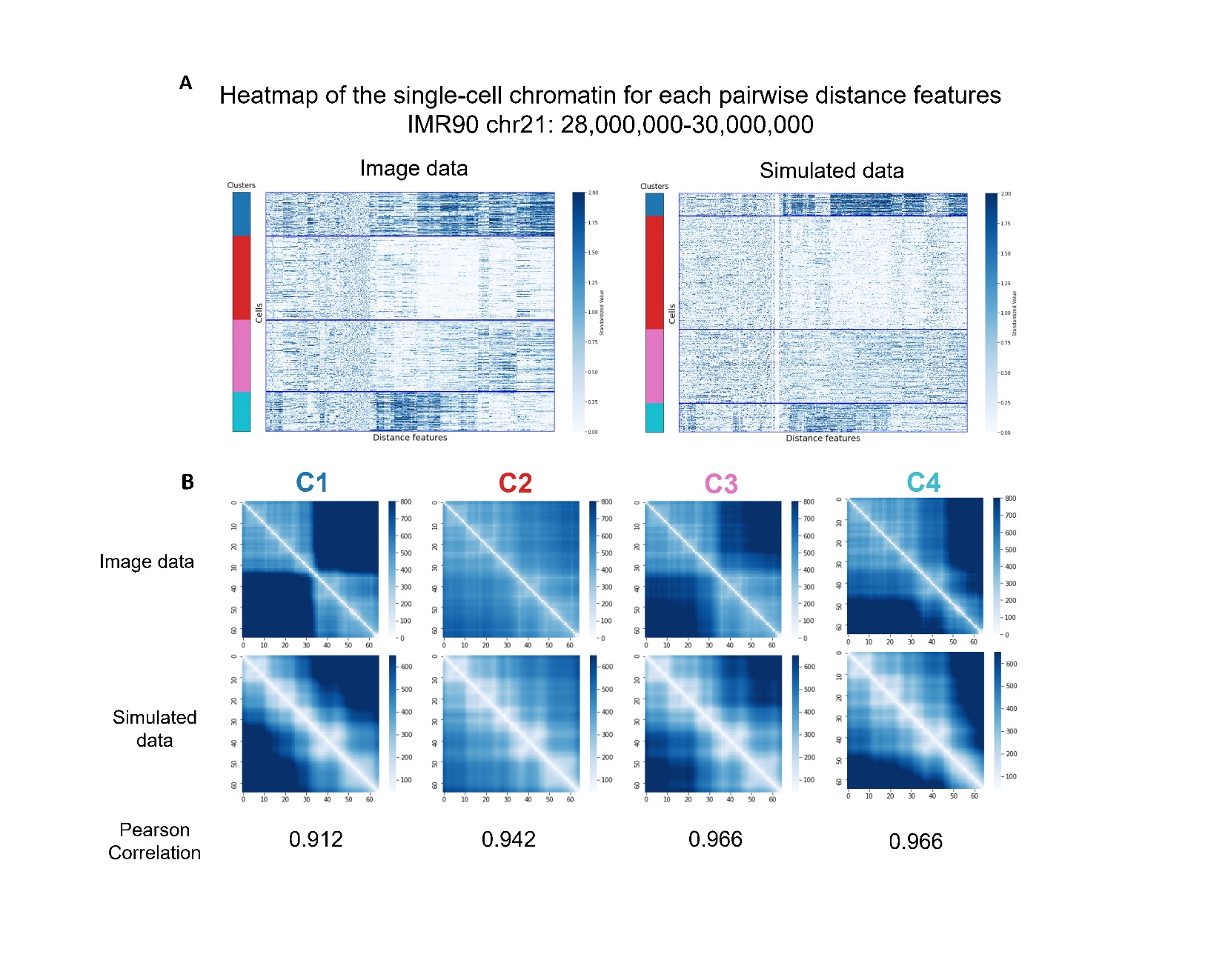

**Supplementary Figure S1**. Comparison of k-means clustering between single-cell chromatin imaging and sBIF-reconstructed structures in IMR90 (chr21:28–30 Mb). (A) Heatmaps of Z-scored pairwise genomic distances for individual cell, with cluster assignments indicated by colored bars (left: imaging data; right: simulated data). Cells and features are ordered by cluster to emphasize subgroup-specific patterns. (B) Cluster-averaged distance matrices for clusters C1–C4, shown for imaging data (top row) and simulated structures (bottom row). Pearson correlation coefficients between imaging- and simulation-derived mean distance matrices are shown below each cluster, indicating strong concordance of structural features between experimental and computational modalities.

Reference:

1. Li,H. (2013) Aligning sequence reads, clone sequences and assembly contigs with BWA-MEM. 10.48550/ARXIV.1303.3997.

2. Open2C, Abdennur,N., Fudenberg,G., Flyamer,I.M., Galitsyna,A.A., Goloborodko,A., Imakaev,M. and Venev,S.V. (2023) Pairtools: from sequencing data to chromosome contacts. 10.1101/2023.02.13.528389.

3. Durand,N.C., Shamim,M.S., Machol,I., Rao,S.S.P., Huntley,M.H., Lander,E.S. and Aiden,E.L. (2016) Juicer Provides a One-Click System for Analyzing Loop-Resolution Hi-C Experiments. *Cell Systems*, **3**, 95–98.

4. Fabian Pedregosa, Gaël Varoquaux, Alexandre Gramfort, Vincent Michel, Bertrand Thirion, Olivier Grisel, Mathieu Blondel, Peter Prettenhofer, Ron Weiss, Vincent Dubourg, Jake Vanderplas, Alexandre Passos, David Cournapeau, Matthieu Brucher, Matthieu Perrot, Édouard Duchesnay; 12(85):2825−2830, 2011.
